# Universal guide for skull extraction and custom-fitting of implants to continuous and discontinuous skulls

**DOI:** 10.1101/2022.01.22.475298

**Authors:** Zurna Ahmed, Naubahar Agha, Attila Trunk, Michael Berger, Alexander Gail

## Abstract

Intracranial neurophysiological recordings require chronic implants to provide transcranial access to the brain. Especially in larger animals, which participate in experiments over extended periods of time, implants should match the skull curvature to promote osseointegration and avoid tissue and bacterial ingress over time. Proposed CAD methods for designing implants to date have focused on naïve animals with continuous and even skull surfaces and calculate Boolean differences between implant and skull surface to fit the implant to the skull curvature. However, custom-fitting by calculating the difference fails, if a discontinuous skull surface needs to be matched. Also, the difference method does not allow designs with constant material thickness along the skull curvature, e.g. to allow fixed screw lengths. We present a universal step-by-step guide for custom-fitting implants which overcomes these limitations. It is suited for unusual skull conditions, like surface discontinuities or irregularities and includes virtual bending as a process to match skull surfaces while maintaining implant thickness. We demonstrate its applicability for a wide spectrum of scenarios, ranging from complex-shaped single-pieced implants to detailed multi-component implant systems built on even or discontinuous skull. The guide uses only a few software tools and the final virtual product can be manufactured using CNC milling or 3D printing. A detailed description of this process is available on GitHub including step-by-step video instructions suitable for users without any prior knowledge in CAD programming. We report the experience with these implants over several years in 11 rhesus monkeys.

**Significance Statement:** Chronic implants are essential for intracranial neurophysiological recordings. In this study we show how to custom-design and –fit such implants for rhesus monkeys (Macacca mulatta). Different to existing approaches, our procedure is not limited to even skull surfaces but can be applied to discontinuous or irregular surfaces. It furthermore presents a description of virtual implant bending to match the skull curvature while maintaining implant thickness. The final virtual product can be manufactured using CNC milling or 3D printing. In contrast to previous studies, this guide is suited for users without any prior expertise in CAD programming using our step-by-step video instructions.

## Introduction

Cranial implants are essential for invasive brain neurophysiology in non-human primates and other animals. For example, headposts are routinely used to stabilize the animal’s head and chamber implants are used to protect craniotomies, which provide access to the brain for intracortical electrophysiological recordings [2,12,13]. With an increasing variety of available neurophysiological recording and stimulation techniques, there is a growing demand to custom-design implants and to custom-fit them to the individual animal. These include advanced chamber designs for semi-chronic adaptive multi-electrode arrays [9, 14, 15] or chronic electrode arrays with wireless recording [3, 28].

Due to the requirements that cranial implants in systems neuroscience are often specific to a recording technique or a project-specific experimental setting, ready-to-go commercial solutions are mostly not available and out-sourcing of the implant design can be expensive and time consuming. Since affordable or even free tools for computer-aided design (CAD) became powerful and production with different materials via 3D printing or CNC milling became easier to access, more and more labs invest into their own CAD-based implant construction.

Previous studies have focused on surgery planning [16, 19] and explain how to extract a 3D skull model from anatomical computer tomography (CT) or magnetic resonance imaging (MRI) scans. The model is then used to physically bend an originally non-molded implant to conform to the skull curvature [18, 24]. Such two-step procedure allows the user to start out with a simpler, non-molded (“standard”) implant which is easier to produce, e.g. on a 3-axes instead of 5-axes milling machine, due to surfaces which only curve along one dimension. Compared to fitting standard implants to the skull during surgery, the existence of a 3D skull model allows shorter surgery times as the fitting is done prior to surgery. Yet, physical fitting might prevent from an optimal fit and increase chances of implant failure due to tissue growth between implant and bone. Also, post-production physical bending can weaken the implant materials and is discouraged for certain metal materials (e.g. titanium) or not possible for plastic materials (e.g. PEEK).

Newer studies have been focusing on methods that allow 3D implant matching to the skull curvature prior to its manufacturing [17] using software like Blender (https://www.blender.org/) or SolidWorks (Dassault Systèmes SolidWorks Corporation, Waltham, Massachusetts) for invasive implants [6,8] and other non-invasive methodologies, e.g. EEG [27]. The resulting implants can be produced using CNC milling or 3D printing. Openness to different production pipelines gives large flexibility in the choice of material. This way, implants can be build that are sturdy, yet small and light-weight, or even radiotranslucent, for compatibility with different imaging techniques.

Previous methods in neuroscience research focused on first-time implantations with smooth skull surfaces that cannot be used for animals with discontinuous skull characteristics [6, 8, 21]. The latter might, for example, result from previous surgical procedures. Studies on human reconstructive surgery, on the other hand, have tried to reconstruct skulls with a hole caused by a previous trauma. Existing approaches mirror the image of the contra-hemispheric skull and calculates the difference between the original skull with hole and the mirror image extracting the missing part of the skull for reconstruction [20, 29]. However, this method assumes perfect symmetry and is not applicable if the skull defects affect both hemispheres.

We present an approach to overcome shortcomings of existing methods. First, our guide enables custom-fitting of implants for animals with unusual skull conditions, such as discontinuities or irregularities on the skull surface. Second, our guide is universal as it is applicable to variable implant designs, including a description of how to virtually bend implants to fit the skull curvature thus allowing to maintain the thickness of the implant even after custom-fitting. Our guide provides a complete process description for customized implant design from skull extraction using imaging data to the final design of the implant in a production-ready file format. Outsourcing of the CAD fitting process to external companies is not necessary while the result of the process can be used for in-house or external production. Final CAD models can be produced by CNC-milling or 3D-printing methods in a large choice of materials. Our extensive tutorial, including step-by-step video tutorials, allows researchers without prior CAD experience to design custom implants. The given examples in this guide are focused on but not limited to non-human primates.

In the following, we will describe how to extract 3D models of the brain and skull followed by the implant design procedure for three categories of implants. To demonstrate the functionality especially for discontinuous skulls, we present its application on a skull containing prior craniotomies by designing a multi-compartment chamber covering the craniotomies. In the end, we will give an overview of the manufacturing processes and file formats.

## Methods

A step-by-step written tutorial guide with corresponding video tutorials for each step, example CAD models and implants can be found on https://github.com/ZuAh/Custom-fitting-of-implants as Extended Data. References to the Extended Data Tutorial will follow the format “ETD 0-1” to indicate chapter (0: “skull extraction”) and processing step (1).

### Skull and brain extraction and locating regions of interest

While the focus of this step-by-step guide is implant design and customization, we will still give a short overview of how to extract a 3D skull and brain model, since this is used as the basis for implant fitting.

We use computer tomography (CT) and magnetic resonance imaging (MRI) to extract the 3D skull and brain models, respectively (Fig. 1). For DICOM image processing we use 3D Slicer (https://www.slicer.org) - an open source software available for Microsoft Windows (Redmond, Washington, USA), Apple Mac OS X (Cupertino, California, USA) and Linux OS. A T2-weighted MRI scan is used to extract the brain (see below). The MRI scan could be used for skull reconstruction as well, but we preferably use CT scans if available due to faster scanning and ease of use in software flow. CT scans have to be aligned with MRI scans, if implant placement depends on the neuroanatomy of the brain. Image alignment can either be done in 3D Slicer using the Transformation module or in a separate CAD program. We used Fusion 360 (Autodesk, San Rafael, California, USA). CT and MRI imaging data types require their own specific extraction steps described in the following.

**Figure 1:**
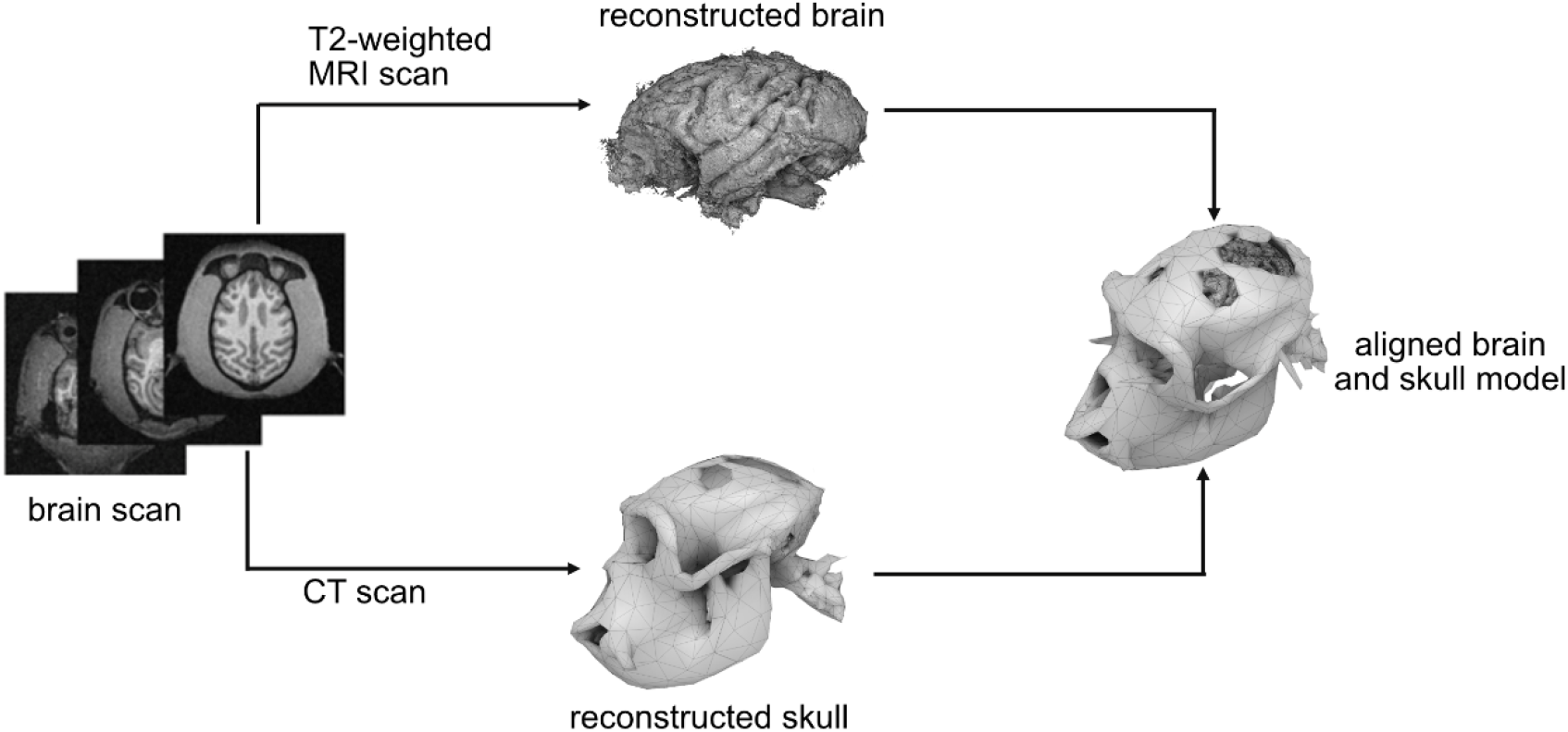
Brain and skull extraction from MRI and CT scans, respectively. Alignment in a stereotactic coordinate frame is achieved using the anatomical landmarks of inferior orbital edge and external auditory bony meatus. An example of a discontinuous skull is shown for demonstration purposes.

Before planning the surgery, the scans are placed in Horsley-Clarke stereotactic coordinates. In 3D Slicer, we used translation and rotation to create a head-centered Horsley-Clarke coordinate system by identifying five points via cranial landmarks: one point in each ear channel (*external auditory bony meatus*), which may be additionally indicated by a fiducial marker from the stereotactic frame during the scan, if a monkey was placed in such; one point below each eye marking the inferior orbital edge (*infraorbital margin*). The fifth point is defined as the equidistant point between the two ear channels and serves as the origin. The scans are translated and rotated until the origin point has the value 0 in a set of three orthogonal coordinate planes and all five stereotactic points are aligned on the horizontal plane.

To extract the skull by using a CT scan, we use the built-in threshold method of 3D Slicer “ThresholdEffect”. This function selects all voxels in the source volume within a range of the signal intensity. The range is modified until the entire skull surface is detected while skull irrelevant voxels are rejected by the algorithm. The 3D skull model can be generated afterwards using the selected voxels (ETD 0-1).

To extract the skull and brain if a T2-weighted MRI scan is available, we use a technique which requires more manual steps. For 3D slicer (version 4.8.1 - S4) the extension named “FastGrowCut” needs to be installed. In newer versions - starting from 4.10.1 - this extension is integrated in the main software and is called “GrowFromSeeds”, which is a multi-label segmentation method. In this approach, seeds are placed by the expert viewer in the region of the skull or brain, respectively. Different seeds can be planted for different tissue types in parallel; for example, one seed type for brain and one for skull. The algorithm detects the skull and the brain separately, such as that one model for each seed type is generated. To verify if the correct areas were detected, it is important to control the tissue separation in the individual slices. If necessary, additional seeds can be placed or suboptimal seeds removed to improve the skull and brain extraction (ETD 0-2).

To plan the coordinates of the implant on the skull if the position of the implant depends on the brain anatomy, we first identify the coordinates of the brain region of interest (ROI). We use anatomical landmarks and a brain atlas [26] to locate the ROI in stereotactic coordinates. We mark the coordinates of ROIs in the 3D skull model by using “MarkUps” and determine the position of the implant on the skull, e.g. by projecting the ROI position in the brain to the surface of the skull along the stereotactic Z-dimension. Another possibility of determining the implant position, especially when targeting deeper brain regions and aiming for surface-normal implant positioning, is the open software package ‘Planner’ [22]. We mark the stereotactic points (origin, ear bars and eye bars) to allow placements of the models in a stereotactic coordinate even after export to a CAD program for implant designing. These marks can additionally be used for alignment of the skull and brain models, if they were obtained from different sources (e.g. a MRI scan for brain extraction, a CT scan for skull extraction). This can be necessary, if the MRI scan is for instance not sufficiently clear to extract the skull model in its details.

We export the extracted 3D brain and skull models in the STereoLitography format (.STL file) (ETD 0-3). This file format can be used for 3D printing of physical brain and skull models, as well for importing to CAD programs for the following skull-fitting implant design procedure (ETD 1-1). Before printing, it is recommended to clean up, and if necessary cut the 3D skull model for better printing quality. We use, for example, the freeware Meshmixer (https://www.meshmixer.com/) for cutting and Meshlab (https://www.meshlab.net/) for quick .STL viewing.

To allow below procedure of implant fitting based on a continuous representation of the skull surface, we create a 3D surface in NURBS (non-uniform rational B-splines) format out of the extracted 3D skull model using Rhinoceros 6 (Robert McNeel and Associates, Seattle, Washington, USA). This is necessary as the STL format of the original imported extracted 3D skull prevents the use of the fitting tools described in this guide.

For this, first, we create a fine mesh using the parameters “Spacing”, which is the space between the individual mesh points, “AutoSpacing”, which enables Rhinoceros to identify the spacing automatically and “AutoDetectMaxDepth”, which detects the depth, can be adjusted (ETD 1-2). A sufficient mesh is created once it covers all relevant parts of the skull surface to a degree of detail that is required for skull reconstruction depending on the *<u>skull condition</u>*. For all our designed implants the automatic “AutoSpacing” and “AutoDetectMaxDepth” with a “Spacing” of 5 was sufficient. Then the mesh is converted in a 3D NURBS surface using “Drape”, which is described in more detail in “*<u>implant design and fitting</u>*” (ETD 1-2).

In the next section, we will describe the implant design processes for this most typical case of a naïve skull being prepared for a first-time implantation. Below, we will return to the topic of skull extraction and reconstruction in the case of more complex surgical situations, e.g. bone discontinuities due to prior surgeries.

### Implant design and fitting

This guide can be used to design various types of implants (Fig. 2). As an example, for a single-compartment implant, we will show how to create a headpost implant (Fig. 2-B). This design is characterized by a central pin (the “post”) extruding vertically from a base. Multiple “legs” build the base and extrude horizontally along the surface of the skull. The legs need to fit the curvature of the skull, but at the same time keep their thickness along the entire length (“virtual bending”), such that bone screws fit exactly the holes in the legs.

**Figure 2:**
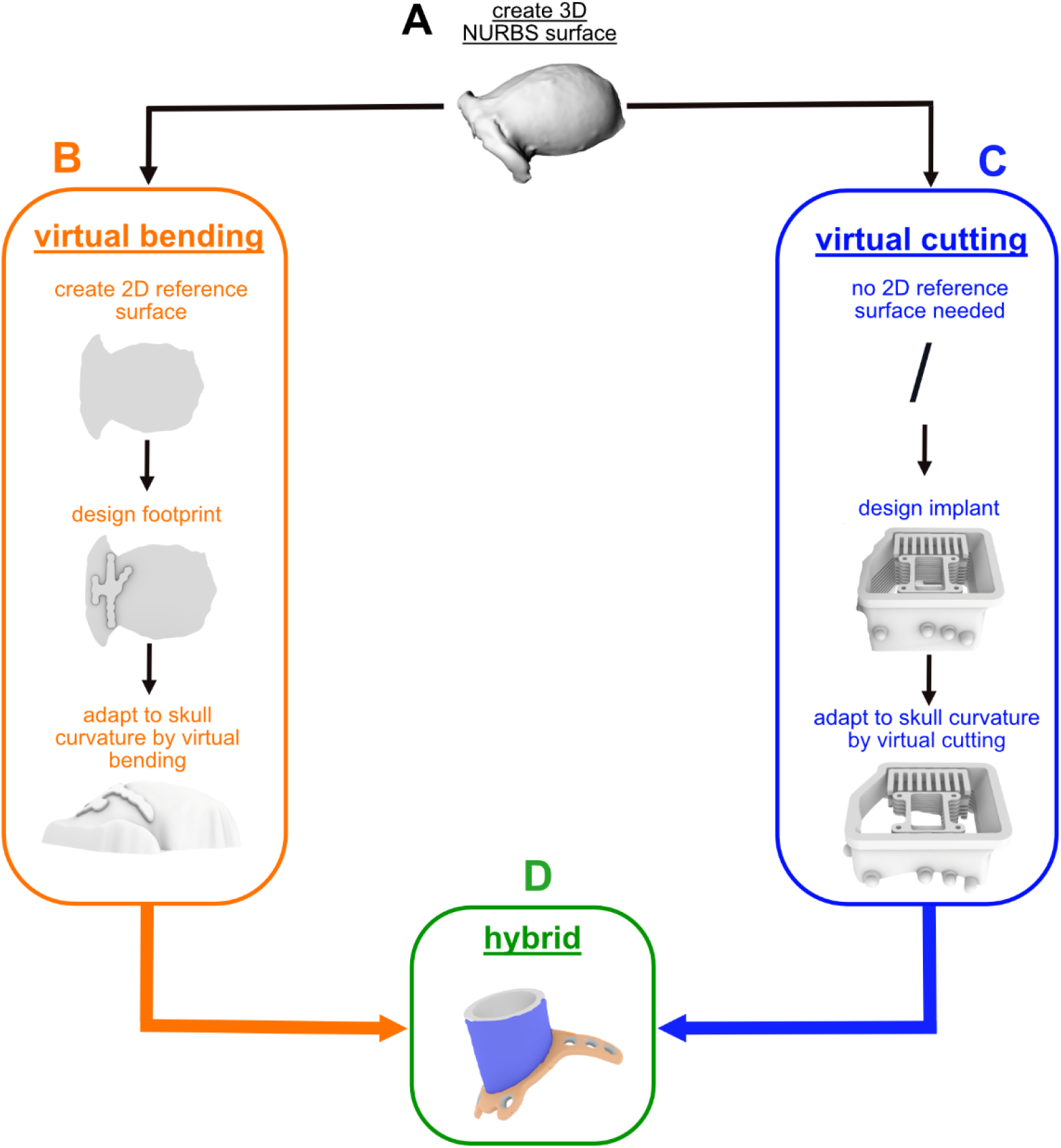
Overview of presented implant types and their individual custom-fitting approaches. A) 3D skull surface in NURBS format reconstructed from the extracted 3D skull surface acquired from a CT scan. B) Virtual bending (orange): After creating a 2D reference surface the to-be-matched implant part is designed and extruded. It is then virtually bent before completion, which implies the thickness of headpost “legs” (perforated metal strips) is maintained while fitting them to skull curvature. C) Virtual cutting (blue): The lower end of large-scale “chamber” (enclosure with lid) is fit to the skull curvature such that remaining height matches desired specification. The example shows a wireless recording chamber with additional interior elements to hold a circuit board and multiple electrode array connectors [3]. D) Hybrid (green): example of a standard chamber to access single brain region of interest (ROI) with legs for mounting; this design combines virtual bending(orange) and cutting(blue).

As an example for a multi-compartment implant, we describe how to create a chamber with different interior elements suitable for chronic array recordings with wireless transmission (Fig. 2-C). One single chamber surrounds and protects a large-scale craniotomy with chronic implanted arrays in multiple brain regions of interest while giving enough space for array connectors and adaptors to hold the wireless transceiver. The elements are molded to the skull surface without constraints by the thickness of the material (“virtual cutting”).

Finally, we describe how to design a standard chamber encircling a small craniotomy and having legs for screwing the implant directly to the skull, similar to the headpost (Fig. 2-D). This demonstrates a implant design which combines cutting and bending (hybrid design).

The focus of the paper is on the digital workflow of designing and custom-fitting the implants to curved surfaces, and will follow below. Since some steps in this guide depend on geometric properties of the implants (e.g. cutting or bending), we first briefly describe the main implant features to give an impression of the scope of implants dealt with.

For implant fitting by virtual bending or cutting we used the software Rhinoceros. For designing of extended chambers, we used Fusion 360, which was due to convenience based on prior user expertise and not a necessity. However, the fitting of all proposed implant types was done using Rhinoceros only.

For animals with pre-existing cranial implants, e.g. a headpost, the persistent implants need to be part of the 3D skull model such that additional implants, e.g. a chamber, can be designed respecting the constraints of the pre-existing implant. Ideally, this planning is done for all implants together based on imaging data recorded prior to first implantation. If this is not possible, and instead the secondary implants need to be added to existing implants based on post-implantation imaging data, artifacts might limit the precision of the skull extraction procedure. On the other hand, changes in skull surface that developed since the first implantation, e.g. bone growth in reaction to the primary implants, can be accounted for in this case, possibly allowing a better fit of the to-be-added implant(s).

### Implants with maintained thickness – virtual bending

Headposts are common in non-human primate research to stabilize the animal’s head position and thereby allow precise measurements of eye movements or applying sensitive neurophysiological probes, e.g. semi-chronic microelectrodes. A common headpost design used in our lab consists of a central transcutaneous post surrounded by four subcutaneous perforated metal strips (“legs”) at its base that are custom-fitted to the skull curvature (Fig. 2-B) for better osseointegration. The implant is fixed to the skull by titanium screws through the holes in the legs, equivalent to titanium strips used in cranioplasty. To maintain the thickness of these legs is important as the self-tapping cortical screws have predefined length and should fit the combined thickness of implant material plus skull to provide implant stability while avoiding transcranial protrusion of the screws.

We achieve constant thickness of skull-fit cranial implants by “virtual bending” (Fig. 2-B). We used the software Rhinoceros for designing the headpost. It offers a useful built-in tool for the bending process. Rhinoceros can also be used for designing the implants itself, which we did in this example.

In the first step, we create a 2D reference plane corresponding to the 3D reconstructed skull, which is converted automatically by Rhinoceros into a 3D NURBS surface format (ETD 1-4 to 1-6). The footprint of the implant is then designed on the virtual 2D reference plane (ETD 1-7). Projecting the outline of the 3D skull model onto this plane helps for planning the layout and leg positions. Once the 2D footprint is finalized, the 2D implant sketch is extruded vertically to create an unmolded 3D version of the implant (ETD 1-7). To remove sharp edges, the function “fillet edges” is used (ETD 1-7).

In the second step, the legs are molded by using the “FlowAlongSrf” - function of Rhinoceros (ETD 8). We selected the extruded footprint as an object to flow along a surface. The previously generated 2D reference plane is used as the “base surface” while the 3D NURBS surface is referred to as the “target surface”.

It is important to select corresponding edges or corners on both surfaces to keep the location of the implant. Additionally, it is helpful to place the 2D surface in front of the 3D NURBS surface, otherwise Rhinoceros can confuse the location and instead of bending on top of the skull try to match it from underneath.

Following, screw holes are added. The diameter of the holes is 3 mm to fit the 2.7 mm cortical screws of 6-8 mm length, which will be used during the surgery to screw the implant to the bone. A counter bore with 45 deg. inclination is added to the holes for later embedding the screw heads (ETD 1-11). It is important not to introduce the screw holes and counter bores before the bending step, as otherwise they will be deformed during the bending due to the non-zero material thickness and thereby would lose their functionality.

In the final step, the transcutaneous post is designed. It consists of an elliptic cylinder which is 14 mm high and has a large diameter of 12.7 mm and a small diameter of 8 mm (exemplary size used in our laboratory). The shape of this post and its top end is adapted to fit the counterpart, which will be specific to the experimental setup. In the example shown here, the top consists of a circular cylinder which has a diameter of 6.8 mm and a height of 6 mm and will later be threaded by hand. To round the post edges on the top and to combine it with the bottom part, the “fillet edges” function is used with a radius of 1 mm.

After fitting, the customized implant is placed on the originally extracted 3D skull to verify the fit, location and angle of the implant (ETD 1-14).

### Extended chambers - virtual cutting

Extended chambers for chronic implantation might protect craniotomies that span several brain regions of interest, and can additionally contain interior constructions for holding electronic equipment for wireless recordings [3]. As an example, we designed a set of multiple components (Fig. 2-C): 1) A biocompatible chamber that encircles the craniotomy and houses array connectors and other components; 2) a connector holder that allows for easy positioning of the electrode array connectors and protecting the connectors and cables against mechanical stress during the surgery; 3) a circuit board holder to attach additional electronic components; 4) Various-sized protective caps covering the chamber while containing different wireless headstages and components.

We use “virtual cutting” for such chamber implants, as they do not need virtually bend legs but still require the bottom part to follow the skull curvature while dimensions and angles towards the top part are preserved. Virtual cutting achieves custom-fitting by calculating the Boolean difference between the implant’s bottom and the skull surface. We exemplify this procedure with a chamber-like implant containing additional interior elements (Fig. 2-C).

In the first step, as above, unmolded 3D versions of all implant components are designed first. We used Fusion 360 for the example presented here, but any other CAD program including Rhinoceros is suited depending on experience, convenience and preference (ETD 2-1). If not originally designed in Rhinoceros, we export all parts that need molding to the skull surface to Rhinoceros using the Initial Graphics Exchange Specification (.IGES) format (ETD 2-4).

In the second step, we import the 3D reference skull surface created in Rhinoceros as described in the section about skull extraction above. The implant components are arranged and placed on the 3D skull surface using the landmarks of the skull. While placing the virtual implant it is pushed through the skull far enough such that the whole lower circumference intersects with the skull and that at the same time the desired implant height remains above skull level. The latter can be measured using either the “Distance” or “Length” command in Rhinoceros (ETD 2-5). For this, the original implant needs to be design high enough; in fact, it might be easiest to design it significantly higher than actually needed, since it will be cut anyway.

In the third step, we use the command “BooleanDifference” and select the implant as the target to subtract from and the 3D NURBS surface as the target to subtract with (ETD 2-6). After fitting, we placed the customized implant on the originally extracted 3D skull to verify the fit, location and height of the implant.

Depending on the way the implant is mounted to the skull, additional design features will be necessary. If the implant is going to be embedded into acrylic dental cement, furrows along the surface help to improve mechanical stability due to the form-fitting of the cement flowing into the furrows (ETD 2-7). If the implant should be directly screwed to the bone, instead, eyelets (single-holed “mini-legs”) can be added to the outer surfaces (ETD 2-2 and 2-3) (Fig. 2-C). When creating such eyelets, it is helpful to place them as close as possible to the expected elevation of the skull curvature. This is achieved by placing the eyelets horizontal midline approximately on the skull surface such that half of the eyelet is sticking inside the skull.

### Hybrid implants – virtual cutting and bending

Standard cylindrical cranial chambers typically encircle small craniotomies, e.g. to target one brain area. We here add custom-fitted legs with constant thickness (equivalent to the ones of the headpost) to the cylindrical center part to demonstrate a hybrid design (Fig. 3-D). The lower end of the cylindrical part follows the skull curvature, while the top of it preserves specific dimensions and angles for mechanical adapters (e.g. to hold recording devices) that should not be distorted by bending.

**Figure 3:**
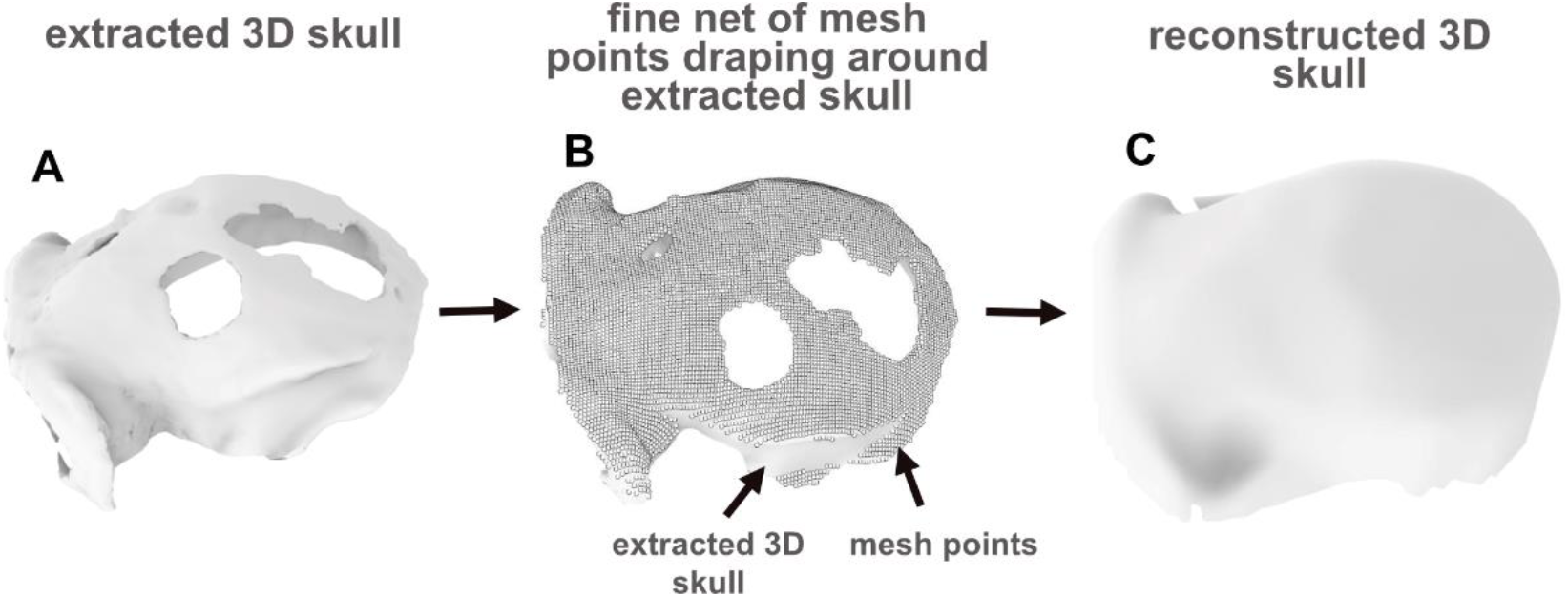
Overview of 3D skull reconstruction for discontinuous or uneven skull surfaces. A) Discontinuous skull with holes reconstructed from CT scan; B) A fine mesh is created out of the originally extracted 3D skull model. All mesh points, which represent the discontinuity, are manually removed. C) Afterwards the mesh is reconverted into a 3D (NURBS) surface.

As a first step, the legs are designed in a process that is equivalent to what is described in the section <u>*Implants with maintained thickness - virtual bending*</u>.

In a second step, the 3D cylinder is designed, in our case with a diameter of 24 mm and height of 30 mm (ETD 3-1). The cylinder is placed on top of the molded legs. We then temporarily remove the legs and push the chamber into the skull model, equivalent to section *Extended chambers* - *virtual cutting*. By calculating the Boolean difference, the cylinder bottom part is fitted to the skull curvature (ETD 3-2). This fitted top part is then placed on the legs and pulled through them until a protrusion of around 1-2 mm on the bottom side of the implant is reached (ETD 3-3). With this (optional) protrusion, the bottom part of the implanted cylinder will be protruding slightly into the craniotomy. Such protrusion of the implant into the craniotomy allows easier centering of the implant on the cranial opening, can help to achieve a better seal between implant and bone and additional mechanical stability, and also can prevent later closure of the craniotomy by bone growth. As the last designing step, the implant is placed on the originally extracted 3D skull to verify fit, location and angle.

### Skull reconstruction based on discontinuous or uneven skull

As key feature, our guide is suitable for designing implants for skulls with preconditions, e.g. discontinuous or uneven skulls. Difficulties in reconstructing 3D skull surfaces with standard techniques can result from unevenness due to excessive bone growth in response to other nearby implants which have been implanted earlier. As part of the osseointegration such bone growth can be stimulated, leading to elevated bone structures on the skull. Skull discontinuities, e.g. left-over skull openings from previous craniotomies, are an additional challenge for implant fitting. Advanced implant fitting techniques should allow, first, discontinuities to be digitally reconstructed and the original shape of the skull to be approximated as best as possible for fitting an implant to areas of the skull that contains bone protrusions or holes with softer tissue. Second, the technique should allow discontinuities to be taken into consideration for designing the implant, e.g. for placing screw holes on top of solid bone structures only and avoiding the open patches. Extra time and effort may be needed to reconstruct the skull and brain when pre-existing implants on the animal’s skull produce scanning artefacts that extend to the part of the skull targeted for implantation.

To virtually reconstruct a discontinuous skull as continuous surface, first, a 2D reference plane is created out of the original 3D surface extracted from the imaging data. This reference plane still contains the discontinuities so that they can be taken into account when designing the implant, e.g. placing screw holes outside the discontinuous regions (ETD 4-6).

In case of an uneven skull (Fig. 3-A), which would prevent reasonably simple implant design (e.g. mandating sharp edges which makes it difficult to manufacture on a milling machine), smoothing out of the unevenness might be required and would later during surgery have to be complemented with corresponding smoothing of the actual bone structure.

Importantly, both the filling of discontinuities and smoothing of unevenness need to be done in a way such that the shape of the other parts of the skull are maintained in their details as much as possible. This is needed to guarantee an appropriate implant match to the actual skull curvature during surgery. We achieve this, after importing the 3D model into Rhinoceros (ETD 1-1), by creating a fine mesh (ETD 1-2) which represents the 3D skull model. We then remove only the single mesh points that are created in the region of the discontinuity or unevenness resulting in a clean mesh point cloud (Fig. 3-B). Finally, we convert the mesh into a 3D NURBS surface by using “MeshPatch” (ETD 1-2) and “Drape” (ETD 1-3; Fig. 3-C).

In contrast to other common smoothing algorithms, our approach allows removing the discontinuity or unevenness in a targeted fashion, i.e. without changing the overall skull surface outside the discontinuous region that would result from general smoothing.

We exemplify the skull reconstruction procedure by designing a headpost and an extended chamber implant for chronic microelectrode implants in an animal with multiple, partly widespread presurgical discontinuities across the skull (Fig. 4).

**Figure 4:**
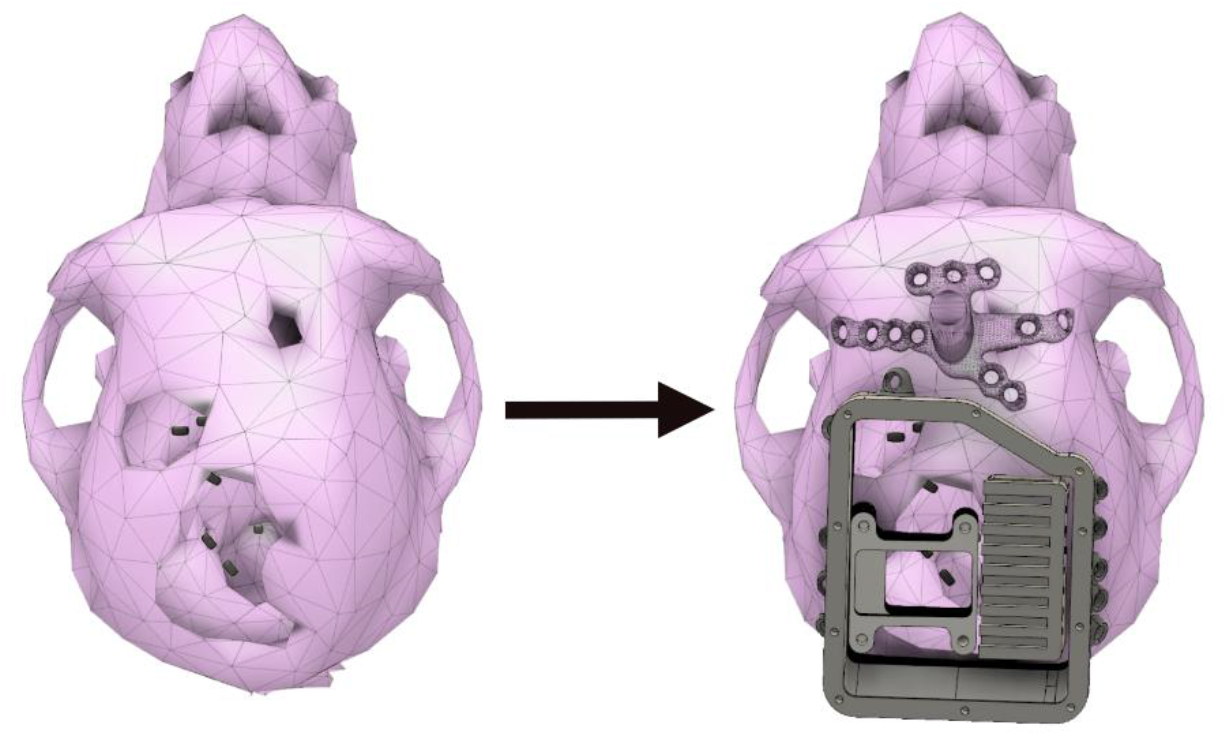
Example of custom-fitted implant for an animal with discontinuous skull surface. Left: The skull contains three holes from previous craniotomies and implants; right: A headpost was designed and custom-fitted taking the anterior hole into account by designing the most posterior leg as a cover for the hole. An extended chamber with inlays for the use with wireless headstages was matched to the skull curvature around two large pre-existing craniotomies. Black rectangles on the cortical surface mark the planned microelectrode array positions.

### Implant manufacturing and file formats

We used in-house 3D printing with Polylectide (PLA) on an additive manufacturing 3D printer (Bibo 2 Maker E, Shaoxing Bibo Automatic Equipment Co., Ltd, Zhejiang, China and Formlabs, Somerville, Massachusetts, USA) for rapid prototyping of all implants and for a physical 3D skull model. Implant dummies together with a printed skull model allow to control for the accuracy of the implant fit after production and to simulate surgical procedures. Prototypes are not necessary for the final implant production. Final implants are milled in-house using a 5-axis CNC milling machine or 3D-printed by commercial services (EOS P770, P396, P110, Shapeways HQ, New York, New York, USA).

For headposts presented here, we used titanium due to its biocompatibility [25], lightweight sturdiness and good chance of osseointegration. It was either milled out of a block of titanium (in-house) or 3D-printed (EOS M280/M290, Sculpteo, Villejuif, France). In case of printing, the thread for the pin necessary for our head-fixation pole was cut afterwards and not included in the print itself as it is too fine to be printed.

Chambers were milled (in-house) out of a block of Polyetheretherketon (PEEK) to provide biocompatibility, MR-transparency and smooth surfaces. During surgery, chambers were screwed onto the skull by using ceramic screws (6-8mm, Thomas Recording,Gießen, Germany) in case of small chambers around a craniotomy or titanium screws (6-8mm, Synthes, West Chester, Pennsylvania, USA) in case of the large chamber. There is no need for applying acrylic dental cement to the bone and implant for mechanical stability with this approach, while cement or biocompatible glue might still be useful in small amounts for sealing the inner side of the chamber against the outside.

Chamber inlays not getting into contact with organic tissue, e.g. because a thin layer of cement covers bony surfaces inside the chamber, were 3D printed using selective laser sintering with lightweight plastics (PA2200, Shapeways) if produced externally; in-house, we used fused deposition modeling with polyactid acid (PLA) on the Bibo printer).

For all 3D printings we use the ‘Stereolithography’ (.STL) data format, while for CNC milling the ‘STandard for Exchange of Product model data’ (.STEP) format was used. In general, for switching between different CAD programs, the. STEP format is recommended as it is supported universally across different CAD programs. However, for importing to Rhinoceros, we recommend to use the ‘Initial Graphics Exchange Specification’ (.IGES) format. This is because. STEP is handled as a block instance in Rhinoceros, which does not allow access to the full range of functions required to custom-fit an implant to the skull.

As 3D slicer uses the. STL format to export the extracted skull and brain models, the original extracted models need to be transformed into NURBS or .IGES surfaces to enable the full functionality of implant fitting tools. After creating a mesh that represents the originally extracted 3D skull model, we converted this mesh into a 3D NURBS surface.

An overview of the used Software Packages can be found in Fig.1-2 in Extended Data.

### Animals and Surgery

Eleven male rhesus monkeys (Macaca mulatta) were implanted with headposts, standard or extended chambers in sterile surgeries under deep anesthesia and peri- and post-surgical analgesia. Data for this study was collected opportunistically, i.e. none of the animals was implanted for the purpose of the current study but instead to be part of neuroscientific research projects. Implant planning was done based on anatomical scans also conducted under anesthesia. Ten of the animals were chronically implanted with a transcutaneous titanium head post. One animal was implanted with a standard chamber with pre-existing headpost. One of the eleven animals was implanted with a chronic chamber for wireless recordings similar to the one described in [3].

All animals were or are housed in social groups with one to two male conspecifics in facilities of the German Primate Center. The facilities provide cage sizes exceeding the requirements by German and European regulations, access to an enriched environment including wooden structures and various toys and enrichment devices [4,7].

All procedures have been approved by the responsible regional government office [Niedersächsisches Landesamt für Verbraucherschutz und Lebensmittelsicherheit (LAVES) under permit numbers 3392 42502-04-13/1100 and 33.19-42502-04-18/2823 and comply with German Law and the European Directive 2010/63/EU regulating use of animals in research.

## Results

### Versatility and efficiency of the design process

Our universal guide describes how to design a variety of implant types and how to deal with special preconditions of the surgical subject, like discontinuous skulls (e.g. skull with holes) and uneven skulls, with a small set of software tools. We so far custom-fitted twelve implants with this approach (table 1): ten headposts (virtually bent), one extended chamber (virtually cut) and one standard chamber (hybrid).

**Table 1:** Overview of implanted animals. Implant types, design process and presurgical skull conditions are indicated. In total 11 animals were implanted with 3 types of implants

|  |  | Implant type | Design process | Skull condition |
| --- | --- | --- | --- | --- |
| 1 | Animal H | extended chamber for wireless recordings & headpost | Virtually cut & virtually bent | Pre-implanted with persisting holes |
| 2 | Animal Z | headpost | Virtually bent | Pre-implanted with persisting holes |
| 3 | Animal T | Standard chamber | Hybrid | Pre-implanted intact |
| 4 | Animal Be | headpost | Virtually bent | Non-implanted intact |
| 5 | Animal Bi | headpost | Virtually bent | Non-implanted intact |
| 6 | Animal J | headpost | Virtually bent | Non-implanted intact |
| 7 | Animal E | headpost | Virtually bent | Non-implanted intact |
| 8 | Animal S | headpost | Virtually bent | Non-implanted intact |
| 9 | Animal D | headpost | Virtually bent | Non-implanted intact |
| 10 | Animal P | headpost | Virtually bent | Non-implanted intact |
| 11 | Animal C | headpost | Virtually bent | Non-implanted intact |

We exemplify the design processes with an extended chamber and its inlays and a headpost, both fitted to a discontinuous skull containing holes (Fig. 4).

The design process is versatile. Besides the custom-fitting of implants by virtual cutting, our guide additionally describes the procedure of virtual bending, which maintains the thickness of an implant when molding it to the skull curvature (Fig. 5-B). All implant types presented here can be customized to both skull conditions (Table 1, Fig 5-A). Within the same framework and with the same set of tools, all implant types can be prepared for production in various metal and non-metal materials using milling or printing methods.

**Figure 5:**
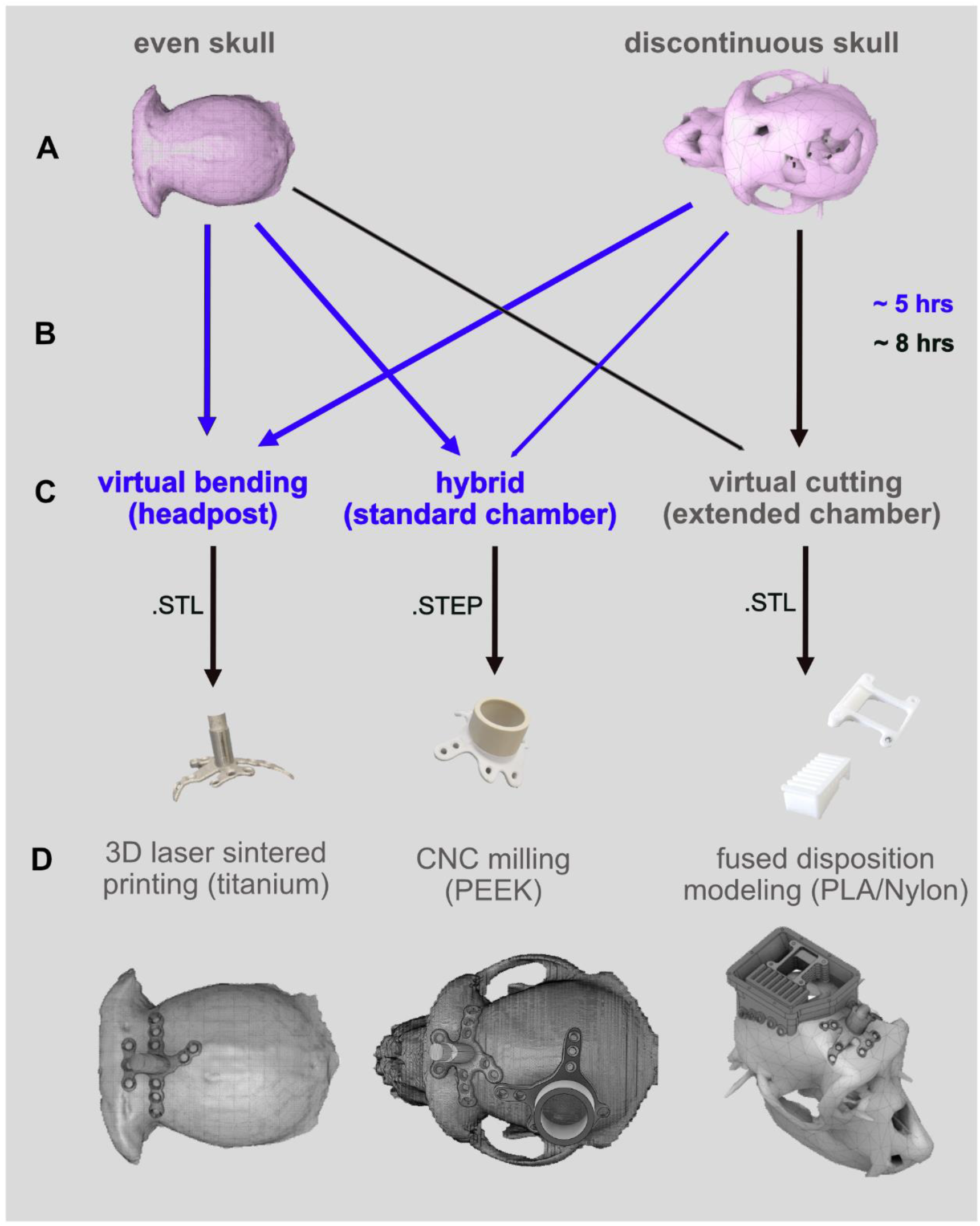
Versatility of our design process. A) Our approach is suitable for intact and discontinuous skulls. B) Arrows indicate the presented combination of skull condition and implant type. Lines indicate possible combination of skull condition and implant type, which were not presented in this paper. The process and guide are adapted to users without any prior knowledge in CAD programming. Single-piece designs were achieved within approximately 5 hours designing time (blue), more complex implant systems within 8 hours (black). C) Three types of implant fitting methods are covered: virtual bending, virtual cutting and the combination of both (hybrid); D) The resulting designs are producible in various (bio-compatible) implant materials.

Our step-by-step guide is efficient and highly accessible. In contrast to previous methods, our proposed way of using the mentioned software packages does not need prior expertise in CAD programming. For users who were naïve to our guide and to CAD programs, reconstruction of a 3D skull model required net five hours and the design of a standard chamber or headpost another five hours (approximate durations). Approximately eight hours were needed to design the extended chamber with its inlays for users who were not familiar with general CAD programming or the procedures described in this guide (Fig.5-B).

### Implant functionality and longevity

With our guide created designs that adhere to the following criteria: 1) manufacturing in the desired biocompatible material, 2) close fit of the implant to the skull without need for further bending or cutting modification during surgery, and 3) functionality of the implant for extended periods of time after implantation.

We consider an implant to be a close-fit, if it sat flush against the skull without demanding any physical modification during surgery. Figure 6 depicts implants, which we classified as “closely fitting implants”.

**Figure 6:**
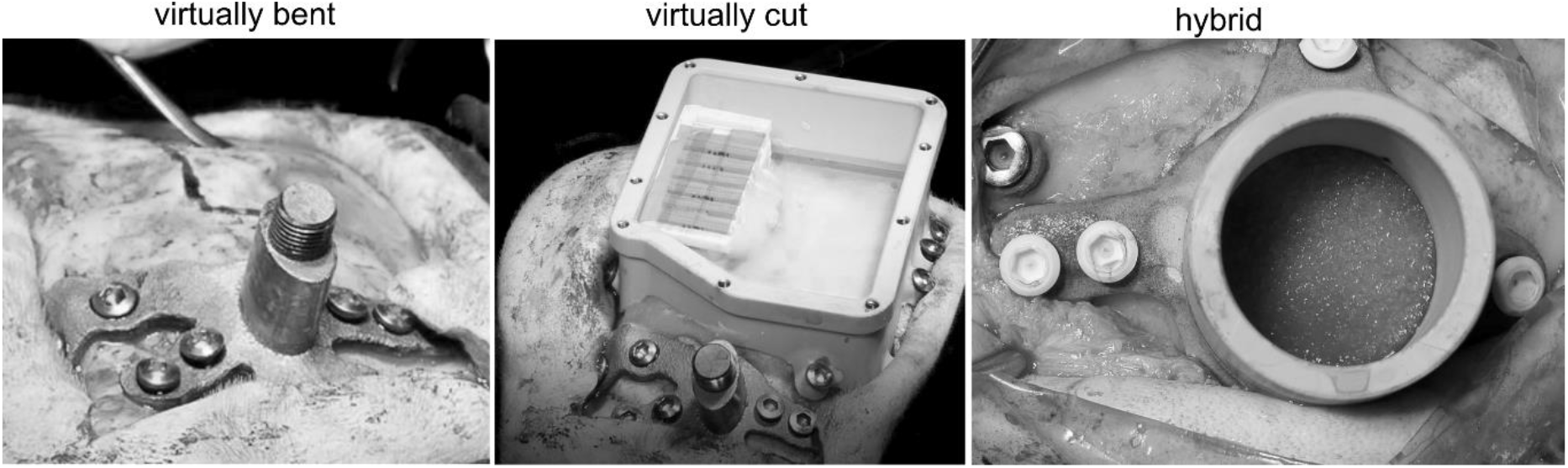
Examples of close-fitting implants. Left: titanium headpost on a discontinuous skull with holes, which was designed by virtually bending; middle: extended chamber for array recordings with its inlay on the same discontinuous skull; right: standard chamber created by virtually bending the legs and virtually cutting the top part (hybrid).

The implants mostly showed lasting functionality. At the time of submission of this manuscript, eight out of ten headposts, which were first-time implantations, are stable and functional up to three years post implantation. Inflammation at the wound margins compromised animal H’s headpost functionality after 4 months; since there was no scientific need use of the headpost was discontinued while use of a second implant on this animal was continued. Also animal Z’s re-implanted headpost required surgical intervention 9 months post-implantation.

The extended chamber for wireless recordings, which is covering two-thirds of animal H’s discontinuous skull is stable up to date (1182 days post-implantation) requiring a small intervention on the chamber inside after 896 days post-implantation. The standard chamber in animal T is robust till date (385 days post-implantation).

Figure 7 shows an example of osseointegration of a titanium headpost produced with a precursor of this guide. As most of the headposts of this paper are still functional, we do not have such documentation yet for these headposts.

**Figure 7:**
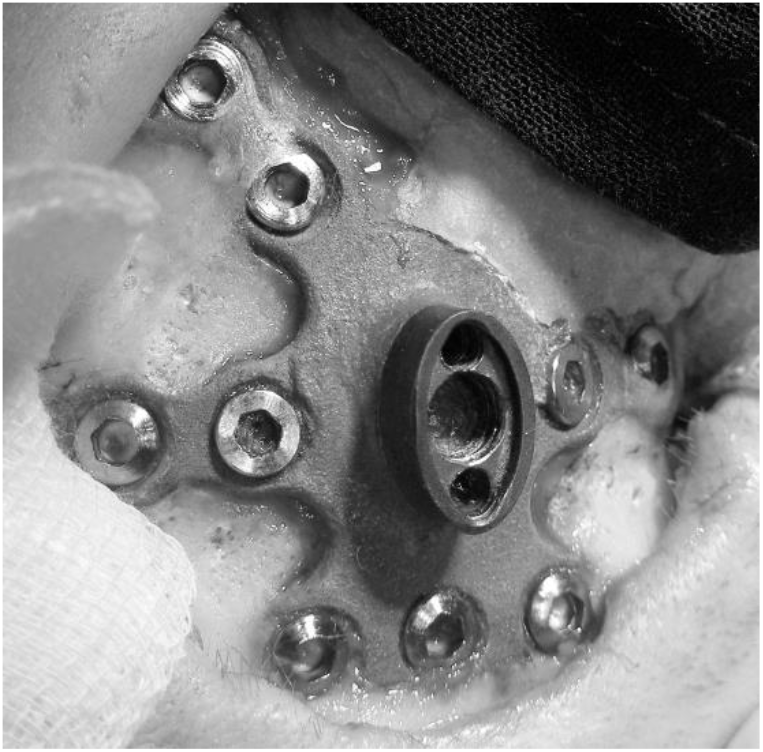
Example image of osseointegration of titanium headpost in bone of the skull.

**Figure 8:**
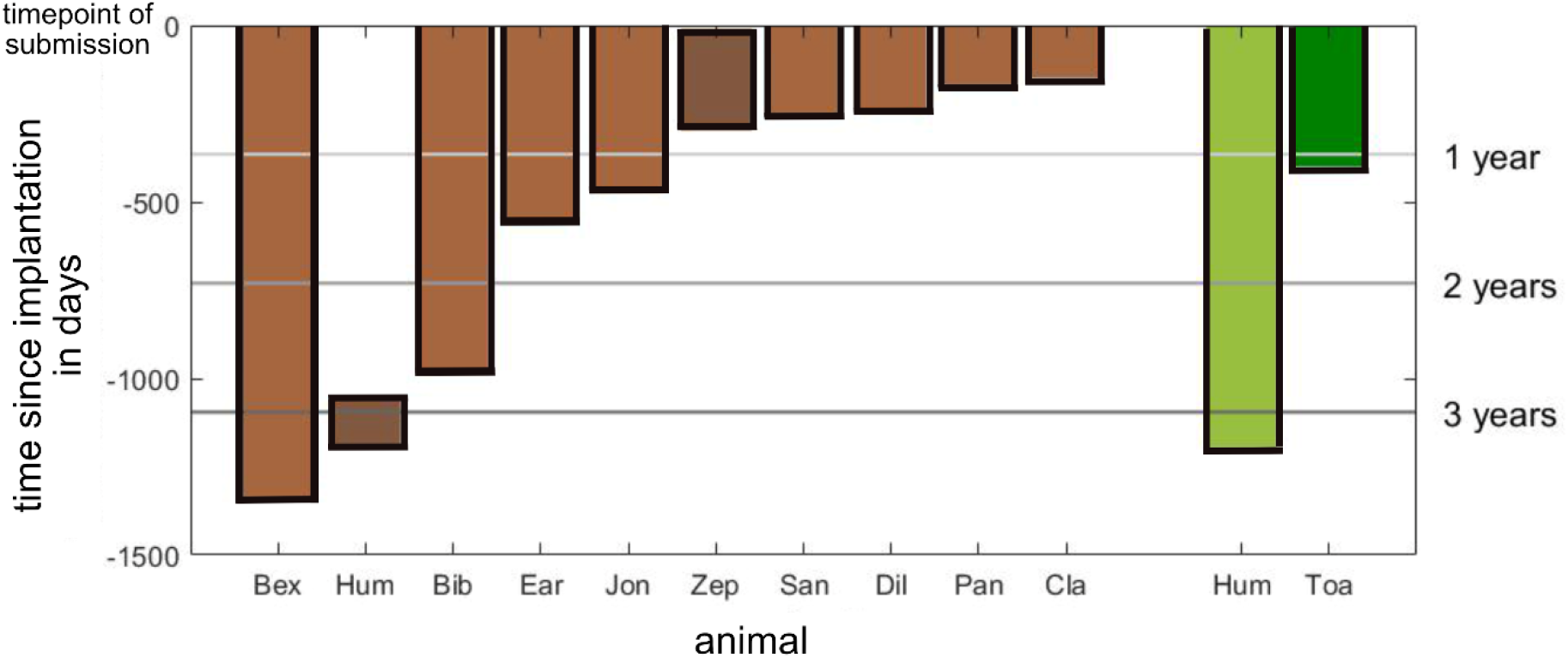
Summary of implant duration in days. Light brown bars indicate days since headpost implantation with still lasting functionality by the time of submission of this manuscript; dark brown: duration of 2 headposts, which lost their functionality; light green: extended chamber implanted on a discontinuous skull; dark green: implant duration of the standard chamber-both still intact.

In summary, using our beginners-friendly guide we designed 12 custom-fit implants of different complexities of which 10 (83%) still remain functional up-to 1329 days (>3 years) post implantation.

## Discussion

We present a universal guide, including video tutorials (in Extended Data), to create implants of diverse types for skulls in various conditions, e.g. smooth skulls and discontinuous or uneven skull surfaces. The guide is well accessible to users who are not CAD experts already. Additionally, to the common method of virtual cutting, we introduce virtual bending and the combination of both. The resulting custom-fitted implant designs and file formats can be used for CNC milling or 3D printing. Using this approach, we successfully implanted eleven animals with headposts, one with a simple cylindrical and one with an extended multi-compartment chamber in animals with even and discontinuous skull surfaces, and demonstrate lasting functionality.

Previous studies have shown that a close implant fit to the skull reduces the risk of sub-implant tissue growth and bacterial infections thereby improving implant longevity [5, 8, 17]. When starting out from non-molded implants with predefined standard shapes that only approximate but do not fit the individual animal, the gap between bone and implant is often minimized by either using acrylic dental cement [8,19] or by physically bending implants to the skull curvature [1,2].

There are disadvantages of acrylic dental cement despite its common use. As acrylic undergoes an exothermic reaction, the heat enhances the risk of bone necrosis [9, 11, 23]. As the cement does not bind to the underlying skull directly, tissue growth can create a gap between implant and bone over time, which may increase the risk of implant instability and infections [1, 5, 24]. Tightly fitting titanium implants, in contrast, can become integrated into the bone, preventing such unwanted tissue growth [19]. Also, thin implant strips on the bone, like the “legs” of our implant designs can be covered with skin. Stabilizing implants by form-fitting with cement requires anchoring screws in the bone. The overall larger volumes of cement which typically cannot be covered with skin, create larger skin openings with more extensive margins that need to be cleaned and protected against infections regularly. Screwing the implant directly to the bone, instead of using anchoring screws plus form-fitting with cement, also allows to adjust the pressure with which the implant presses against the bone when tightening the screws.

As an alternative, non-molded implants, which are commercially available or produced in a standard shape without custom-fitting to the skull, could be used and be physically adjusted during surgery or prior to surgery when a 3D printed skull replica is available as template. This method is limited by material constrains, though. Plastic materials mostly cannot be adjusted in shape by bending but only by cutting or filing off. Metal materials instead are difficult to cut or file off especially during surgery. Brittleness and reduced durability after hot-bending or forging prohibits larger changes in shape, for example for titanium implants [1,18, 22, 24]. Physical cutting, filing or bending may lead to imperfect skull fitting, making the osseointegration more challenging as it can result in gaps between implant and bone. For large-scale implants precise fitting and the associated challenges are particularly relevant [6].

More recent work suggests designing custom-fitted implants in 3D CAD programs ensuring a closer fit to the skull curvature compared to bending of commercially available standard implants [6, 8, 17, 21]. The proposed CAD-based 3D fitting procedures can successfully customize implant shapes, but commonly use the Boolean Difference method (“virtual cutting”) to fit the skull curvature. Constant thickness of implants along the skull surface is difficult to achieve with cutting methods compared to bending. Virtual bending easily allows for thin implant structures to fit screws of constant length and with surface-normal orientation to both bone and implant for optimal direction of the forces, while still being covered by skin. By combining virtual cutting with virtual bending, as described in our guide, the design of a broad range of implants is possible, e.g. single piece headposts maintaining their leg thickness, extended, complex multi-compartment implants or standard chambers with legs, etc.

While previous methods require a certain proficiency in CAD programming, our aim was to provide a method that can be easily learned and used by non-proficient CAD users. Testing with users who were previously naïve about CAD design showed that the guide with its tutorials allows designing of implants of decent complexity and custom-fit to individual animals within a few hours. The final virtual product can then be manufactured using both production pathways, CNC milling or 3D printing.

With our approach, we also wanted to target the challenge of custom-fitting implants for uneven animal skulls, especially skulls with discontinuities, e.g. holes. While such skull conditions probably are not encountered often, the possibility to re-implant otherwise healthy animals and thereby reduce the number of animals needed for a study, is especially relevant for large and long-living animal models, like non-human primates. These animals often underwent very time-consuming preparation for a study, e.g. behavioral training, additionally increasing their value. Approaches in human reconstructive surgery try to virtually reconstruct skulls with holes by producing a mirror image of the contra-lesional half of the skull [20, 29]. However, for this approach at least one unaffected hemisphere is necessary. We demonstrated successful application of our approach in an example for which skull discontinuities spread across both hemispheres. Our guide enabled us to design a headpost, an implant type matched to the skull by virtual bending while maintaining its thickness, and an extended chamber with its inlays for wireless recordings, which was fitted by virtual cutting, to fit the discontinuous skull. It allowed us to customize and plan the implants by including the skull discontinuities without removing other non-targeted uneven features of the skull as it is the case with common smoothing tools. The extended chamber is to date stable, well-integrated and functional.

### Further method refinement

Our guide makes use of different software packages of which only part of their functionality is used. Additional features might help optimizing the planning and implant design. For example, 3D slicer can not only be used to extract a 3D skull and brain model but also to extract vascularity of the animal’s brain (Fig. 1-1 in Extended Data). This can help to improve the surgery planning by avoiding major vasculature, which otherwise might complicate the access to the planned ROI during surgery. A contrasted MRI (e.g. using Gadovist) is necessary to make the blood vessels visible.

The presented and extracted 3D brain models in this guide show a less well defined structure in the anterior brain regions. In our MRI scans, the distinction between brain and non-brain tissue in this area is less distinct and clear as for the posterior part. The brain extraction can be improved by adding seeds for non-brain tissue in the respective regions. As we did not target these anterior brain regions with our chamber implants, we did not invest time to optimize this aspect.

We attempted to use 3D printing techniques for extended chambers out of PEEK. We considered the resulting printed versions of the chamber not sufficiently smooth on the outside surfaces, even after several revisions. Also, the eyelets for screwing the implant to the bone could not be printed well enough. The same file could be printed in PLA during rapid prototyping without these problems, indicating that the yet not very common PEEK 3D printing technique itself might not be sufficient. However, this might become a valid option in the future. CNC milling did not cause any problems and the implants could be manufactured appropriately.

## Acknowledgements

We thank Klaus Heisig and Sina Plümer for helpful discussions with implant design and technical support.

